# Novel full-length major histocompatibility complex class I allele discovery and haplotype definition in pig-tailed macaques

**DOI:** 10.1101/186494

**Authors:** Matthew R. Semler, Roger W. Wiseman, Julie A. Karl, Michael E. Graham, Samantha M. Gieger, David H. O’Connor

## Abstract

Pig-tailed macaques (*Macaca nemestrina, Mane*) are important models for human immunodeficiency virus (HIV) studies. Their infectability with minimally modified HIV makes them a uniquely valuable animal model to mimic human infection with HIV and progression to acquired immunodeficiency syndrome (AIDS). However, variation in the pig-tailed macaque major histocompatibility complex (MHC) and the impact of individual transcripts on the pathogenesis of HIV and other infectious diseases is understudied compared to rhesus and cynomolgus macaques. In this study, we used Pacific Biosciences single-molecule real-time circular consensus sequencing to describe full-length MHC class I (MHC-I) transcripts for 194 pig-tailed macaques from three breeding centers. We then used the full-length sequences to infer *Mane-A* and *Mane-B* haplotypes containing groups of MHC-I transcripts that co-segregate due to physical linkage. In total, we characterized full-length open reading frames (ORFs) for 313 *Mane-A*, *Mane-B*, and *Mane-I* sequences that defined 86 *Mane-A* and 106 *Mane-B* MHC-I haplotypes. Pacific Biosciences technology allows us to resolve these *Mane-A* and *Mane-B* haplotypes to the level of synonymous allelic variants. The newly defined haplotypes and transcript sequences containing full-length ORFs provide an important resource for infectious disease researchers as certain MHC haplotypes have been shown to provide exceptional control of simian immunodeficiency virus (SIV) replication and prevention of AIDS-like disease in nonhuman primates. The increased allelic resolution provided by Pacific Biosciences sequencing also benefits transplant research by allowing researchers to more specifically match haplotypes between donors and recipients to the level of nonsynonymous allelic variation, thus reducing the risk of graft-versus-host disease.

## Introduction

Nonhuman primates (NHPs) are of particular importance for their use as animal models for the study of human immunodeficiency virus (HIV). NHPs infected with simian immunodeficiency virus (SIV) mount similar immune responses as humans infected with HIV (Baroncelli et al. 2008; Gardner and Luciw 2008; Joag et al. 1997). Major histocompatibility complex (MHC) class I proteins play a particularly important role in HIV/SIV immune containment by presenting peptides on the surface of virally-infected cells to CD8+ killer T-cells (Bontrop 2006). It is therefore important to understand the diverse MHC class I (MHC-I) transcript sequences within the population of NHPs that differ in their peptide binding specificity. This can be difficult because many commonly used NHPs, including all macaques, have a complex MHC-I region that can contain over 20 MHC-I genes on each chromosome (Daza-Vamenta et al. 2004; Wiseman et al. 2013).

For many years, rhesus macaques (*Macaca mulatta, Mamu*) have been the preferred NHP model for HIV vaccine development due in part to their well-characterized MHC-I genes. However, the supply of rhesus macaques is becoming constrained due to their popularity (Baroncelli et al. 2008). Cynomolgus macaques (*Macaca fascicularis, Mafa*) are also widely used for SIV research, especially in European countries. Although cynomolgus macaques are an excellent model to study viral replication, they tend to control viremia shortly after initial infection with commonly used SIV strains (many of which were initially adapted to rhesus macaques). Frequent spontaneous control makes cynomolgus macaques suboptimal for most models of HIV vaccine development (Baroncelli et al. 2008; Reimann et al. 2005). Pig-tailed macaques (*Macaca nemestrina, Mane*) are emerging as an important model for HIV/AIDS research and vaccine development. Pig-tailed macaques, like rhesus and cynomolgus macaques, can be infected with SIV and simian/human immunodeficiency virus (SHIV) while also progressing to simian AIDS (Hatziioannou et al. 2009, 2014; Joag et al. 1997). Moreover, pig-tailed macaques, unlike rhesus or cynomolgus macaques, can also be infected, and become symptomatic, when challenged with minimally modified strains of HIV (Del Prete et al. 2016; Frumkin et al. 1995; Hatziioannou et al. 2009, 2014). Infection with a short transcript version of HIV type 1 (HIV-1 ST) progressed to AIDS-like disease in pig-tailed macaques after depletion of CD8+ cytotoxic T cells (Hatziioannou et al. 2014). Following serial passage, HIV-1 ST acquired the ability to antagonize multiple macaque restriction factors, replicate at substantially high levels, and deplete CD4+ helper T cells in a way similar to human progression to AIDSlike diseases (Hatziioannou et al. 2014).

Additionally, infection of pig-tailed macaques with replication competent HIV type 2 (HIV-2) led to a decline of CD4+ helper T cells, a characteristic sign of AIDS-like diseases (Baroncelli et al. 2008). This is likely due to the fact that pig-tailed macaques contain a variant of tripartite motif-containing protein 5 alpha (TRIM5α) differing in several amino acids from the versions found in other macaque species. Normal functioning TRIM5α binds to HIV capsid proteins, thus preventing uncoating and successful replication of the virus inside viable host cells (Kirmaier et al. 2010). The pig-tailed macaque TRIM5α variant binds less tightly to these proteins, leading to progression of HIV infection and release of replication competent virus from host cells (Brennan et al. 2007; Igarashi et al. 2007; Stremlau et al. 2004). Because of this modified protein, HIV-2 infection in pig-tailed macaques has become another important model for studying human infection with HIV and progression to AIDS-like disease (Baroncelli et al. 2008, Hatziioannou et al. 2014).

Even with multiple models to mimic HIV infection, relatively little is known about pig-tailed macaque MHC-I genetics. Previous studies have identified multiple novel haplotypes and allelic variants in different cohorts (Fernandez et al. 2011; O’Leary et al. 2009; Pratt et al. 2006). However, these studies largely used short genotyping amplicons to describe haplotypes and characterize novel MHC alleles - a 195 base pair (bp) amplicon encoding the highly polymorphic region of exon two (Wiseman et al. 2009), a 367 bp amplicon encompassing exons two and three (O’Leary et al. 2009), and a 568 bp amplicon spanning from exon two into exon four (Fernandez et al. 2011). These MHC-I fragments can be useful for genotyping, but are not as informative as full-length MHC-I transcript sequences that can now be easily recovered using Pacific Biosciences (PacBio) circular consensus sequencing (CCS) technology (Karl et al. 2017; Pratt et al. 2006; Westbrook et al. 2015).

In this study, we used PacBio single-molecule real-time (SMRT) CCS technology for full-length allele discovery and haplotype definition in 194 pig-tailed macaques originating from three different institutions. We used primers to amplify the full-length ~1.1 kilobase (kb) sequences that encode the MHC-I proteins. Amplicons were created and amplified from complementary DNA (cDNA), thus allowing us to specifically amplify full-length, functional open reading frame (ORF) transcripts. We defined 236 novel MHC-I transcript sequences and extended 87 previously described MHC-I transcript sequences to include full-length ORF sequences. With the addition of these novel sequences, we expanded the known diversity of MHC-I in pig-tailed macaques, including allelic variants of published sequences that have previously been shown to be protective against SIV and other infectious diseases in both pig-tailed and rhesus macaques (Gooneratne et al. 2014; Loffredo et al. 2007, 2009; O’Connor et al. 2003; Pratt et al. 2006; Smith et al. 2005a, b). We also defined 192 high-resolution *Mane* MHC-I haplotypes which are useful in showing the inheritance of parental chromosomes and also reducing the risk of graft-versus-host disease (GvHD) in transplant studies using MHC identical animals.

## Materials and Methods

### Animal selection for full-length allele discovery

Cellular RNA was obtained from 79 pig-tailed macaques from investigators at Johns Hopkins University (JHU; Baltimore, MD). cDNA was provided from 91 pig-tailed macaques housed at the Washington National Primate Research Center (WaNPRC; Seattle, WA) and 90 pig-tailed macaques from investigators at the University of Melbourne (Melbourne, Victoria, Australia) and the Monash University Animal Research Platform (Melbourne, Victoria, Australia). All animals were cared for according to the regulations and guidelines of the Institutional Care and Use Committee at their respective institutions.

Illumina MiSeq (San Diego, CA, USA) sequencing was performed for all three cohorts as previously described (Karl et al. 2014, 2017) in order to inform selection of animals for PacBio CCS technology (data not shown). Out of the original 260 animals, 194 were selected for full-length MHC-I sequencing. Of these 194 samples, 63 were from JHU, 43 were from the WaNPRC, and the final 88 samples were from the University of Melbourne and Monash University. Samples were selected for PacBio sequencing based upon whether they appeared to carry a novel haplotype(s) or contained a high percentage of reads not mapping to known sequences according to the Illumina MiSeq genotyping results. In order to minimize the amount of redundancy in novel allele discovery, a subset of individuals were excluded since they only carried MHC haplotypes that were shared with multiple other members of the same breeding center cohort. Animals were divided into eight separate pools depending on their institution of origin. These pools had a range of 15-64 animals per pool and each pool was run on between four and eight SMRT cells depending on the number of animals in each pool.

### PCR amplification for PacBio RS II sequencing

We synthesized cDNA from RNA received from the various institutions using Superscript^TM^ III First-Strand Synthesis System for reverse transcription polymerase chain reaction (RT-PCR) (Invitrogen, Carlsbad, CA, USA) on the samples selected for full-length sequencing. cDNAs for full-length MHC-I sequencing were amplified using a combination of two forward and three reverse primers that annealed to the 5’ and 3’ UTRs, respectively. Each amplified product also contained a unique PacBio 16 bp barcode (Menlo Park, CA, USA) that was fused to the 5’ end of the sequence-specific oligos (Supplemental Figure 1). Amplification was performed on an Applied Biosystems Veriti^TM^ Thermal Cycler (ThermoFischer Scientific, Foster City, CA, USA) under the following conditions: initial denaturation at 98°C for 3 minutes; 25 cycles of 98°C for 5 seconds, 60°C for 10 seconds, 72°C for 20 seconds for amplification; and a final elongation of 72°C for 5 minutes before being held at 4°C until proceeding. The full-length products were confirmed on the FlashGel DNA cassette system (Lonza, Basel, Switzerland). After confirmation on the FlashGel, the full-length products were initially purified using the AMPure XP PCR purification kit (Agencourt Bioscience Corporation, Beverly, MA, USA) at a DNA to bead ratio of 1:1. Quantification was performed on the purified products using the Quant-iT dsDNA HS Assay kit and a Qubit fluorometer (Invitrogen, Carlsbad, CA, USA) using a DNA to buffer ratio of 2:198. The final purification was performed using AMPure PB beads (Agencourt Bioscience Corporation, Beverly, MA, USA) at a DNA to bead ratio of 1:1 and quantified using the Qubit fluorometer following the same protocol described above.

SMRTbell libraries were created using the PacBio Amplicon Template preparation protocol for CCS using P6-C4 chemistry where individual molecules are sequenced multiple times in both orientations (www.pacb.com). This protocol was described in detail previously (Karl et al. 2017; Prall et al. 2017). Briefly, the pools of PCR products were repaired and hairpin adapters were incorporated onto the ends of the products using the PacBio DNA Template Prep Kit 2.0. The products were the purified using an AMPure PB beads to DNA ratio of 0.6:1. Quality of the pool was assessed again using the Qubit dsDNA BR assay (Invitrogen, Carlsbad, CA, USA) and the Agilent 2100 Bioanalyzer DNA12000 kits (Life Technologies, Madison, WI, USA) following the manufacturer’s protocol. Full-length amplicons were sequenced on a PacBio RS II instrument (Menlo Park, CA, USA) as was previously described (Karl et al. 2017; Westbrook et al. 2015). Sequencing was performed by the University of Washington PacBio Sequencing Services core or at the Great Lakes Genomics Center at the University of Wisconsin-Milwaukee.

### Analysis of MHC-I full-length sequences

Results of full-length sequences were analyzed as previously described (Karl et al. 2017; Prall et al. 2017). In short, we removed sequenced reads that perfectly matched to previously described full-length pig-tailed macaque alleles that were described using Sanger sequencing along with pyrosequencing. These sequences were available to us in the Immuno Polymorphism Database for the Major Histocompatibility Complex genes of Non-Human Primate (IPD-MHC NHP) (http://www.ebi.ac.uk/ipd/mhc/nhp/index.html). We excluded splice variants and chimeric sequences to leave only valid, full-length novel sequences or full-length extensions of previously described transcript sequences in the IPD. Remaining processed reads were clustered, and clusters containing three or more reads were mapped to closest related previously described sequences. Novel sequences and extensions of previously described alleles were analyzed and validated using Geneious Pro (version 9.0) (Biomatters Limited, Auckland, New Zealand) and Basic Local Alignment Search Tool (BLAST). The BLAST results between the novel and known sequences were compared and the putative novel sequences were given local names with the closest related allele and the number of nucleotides by which they differed. A genotyping table (Supplemental Figure 2) was generated with the number of identical reads identified per candidate allele in each sample. Novel sequences confirmed by this approach as well as extensions of previously described partial sequences were submitted to IPD-MHC NHP to obtain official transcript nomenclature (Maccari et al. 2017; Robinson et al. 2013).

## Results and Discussion

### Full-length MHC-I allele discovery

Our initial pilot study involved a subset of 16 animals from JHU in which identified *Mane* transcript sequences that perfectly mapped to known sequences were not removed in order to provide proof of concept. In total, we identified 35 previously described sequences, along with 39 extensions of previously reported partial sequences and 70 novel full-length ORF transcript sequences that differed by one or more nucleotides compared to their closest related known sequences. This study demonstrated to us that our analytical methods could successfully identify full-length ORF transcript sequences previously characterized by cDNA cloning and Sanger sequencing or by Roche/454 pyrosequencing. In our subsequent studies, all sequences that perfectly matched to previously described full-length *Mane* sequences were removed using our novel allele discovery pipeline and were not included when describing the total number of transcripts discovered. These known *Mane* transcripts were taken into account when describing new haplotypes or updating sequences associated with previously described haplotypes.

As a result of all sequencing runs, each sample across the three facilities had an average of 1,225 reads identified from PacBio sequencing. An average of 15 transcripts were identified for each animal, including novel sequences without formal names at the time. Using PacBio CCS sequencing, we characterized 313 full-length novel ORF sequences and extensions of previously known *Mane* MHC-I sequences as shown in Table 1, as well as the 35 previously described alleles from our initial pilot study. Of these 313 newly characterized transcript sequences, 116 were *Mane-A* sequences, 161 were *Mane-B* sequences, and 36 were *Mane-I* sequences. This distribution matches what we have seen in previous studies of pig-tailed macaques as *Mane-B* sequences are more numerous than either *Mane-A* or *Mane-I* sequences (O’Leary et al. 2009). Of these sequences, 241 were completely novel sequences and 72 were full-length extensions of previously described alleles. We did not identify any *Mane-E* transcripts because the UTR in *Mane-E* transcripts differs significantly from the UTR sequences of *Mane-A, Mane-B*, and *Mane-I* regions. Thus, the primers that we use to amplify *Mane-A, Mane-B*, and *Mane-I* sequences fail to amplify *Mane-E* sequences efficiently.

**Table 1:**
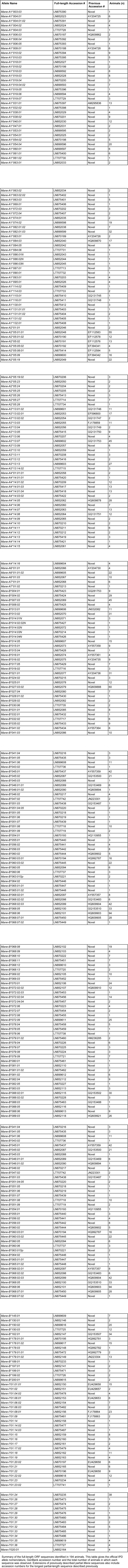
Full-length Mane transcripts characterized by PacBio sequencing

We also discovered two novel variants of *Mane-A1*084:01* and a full-length extension of *Mane-A1*084:03* which was previously described (Fernandez et al. 2011). *Mane-A1*084*, previously named *Mane-A*10*, has been associated with delayed progression of SIV (Pratt et al. 2006; Smith et al. 2005a, b) and is therefore important to investigators studying SIV. The additional variants of *Mane-A1*084:01* may provide more insight to interactions among MHC-I and HIV proteins for specific vaccine design. A single novel variant of *Mane-B*008:01*, and one novel variant and one full-length extension of *Mane-B*017:01* were also discovered as a result of this study. In rhesus macaques, *Mamu-B*008:01* and *Mamu-B*017:01:01* transcript sequences have been associated with exceptional control of SIV replication during the chronic phase of infection (Gooneratne et al. 2014; Loffredo et al. 2007; Martins et al. 2015; O’Connor et al. 2003; Yant et al. 2006), and further studies are necessary to determine if these newly characterized allelic variants show the same protective effect as their *Mamu* counterparts.

We also compared the similarity of our newly characterized alleles to previously described alleles from rhesus and cynomolgus macaques as there is data to suggest commonly shared transcript sequences between the three species (Karl et al. 2017). We identified 37 out of 313 full-length ORF transcript sequences that were identical to previously characterized rhesus or cynomolgus nucleotide sequences, with thirteen mapping to both a rhesus and cynomolgus sequence (Table 2). These shared sequences likely represent diverse MHC-I alleles that were present in a common ancestor of these three macaque species. With increased feasibility of performing high-throughput full-length ORF sequencing on large macaque cohorts, it is likely that more transcripts will appear to be shared between the same species of macaque in different geographical locations and between different macaque species.

**Table 2.** *Mane* transcripts that are identical to previously described Mamu or Mafa class I sequences

| ***Mane* Allele** | **IPD Accession #** | **Identical *Mamu* and/or *Mafa* sequence(s)** |
| --- | --- | --- |
| *Mane-A1\*004:01:01* | IPD0006235 | *Mamu-A1\*004:02:01* (IPD0001811 - 1098 bp) |
| *Mane-A1\*004:01:02* | IPD0007156 | *Mamu-A1\*004:02:02* (IPD0001693 - 1098 bp) |
| *Mane-A1\*032:02* | IPD0007162 | *Mafa-A1\*032:04* (IPD0002384 - 1003 bp) |
| *Mane-A2\*05:19:01* | IPD0006268 | *Mamu-A2\*05:04:03* (IPD0001911 - 1098 bp),<br>*Mafa-A2\*05:06:02* (IPD0001218 - 1098 bp) |
| *Mane-A2\*05:25* | IPD0007191 | *Mamu-A2\*05:10* (IPD0001871 - 1098 bp),<br>*Mafa-A2\*05:57:01* (IPD0006411 - 1098 bp) |
| *Mane-A3\*13:04* | IPD0004048 | *Mafa-A3\*13:17:01* (IPD0006415 - 822 bp) |
| *Mane-A4\*01:01:01* | IPD0004054 | *Mamu-A4\*01:02:01* (IPD0001930 - 862 bp),<br>*Mafa-A4\*01:09* (IPD0003394 - 1074 bp) |
| *Mane-A4\*14:01:03* | IPD0007183 | *Mafa-A4\*14:01* (IPD0001798 - 1098 bp),<br>*Mamu-A4\*14:04:02* (IPD0006920 - 692 bp) |
| *Mane-A4\*14:06* | IPD0007050 | *Mamu-A4\*14:04:02* (IPD0006920 - 692 bp) |
| *Mane-A4\*14:15* | IPD0007051 | *Mamu-A4\*14:04:02* (IPD0006920 - 692 bp) |
| *Mane-A4\*14:16* | IPD0007298 | *Mamu-A4\*14:04:02* (IPD0006920 - 692 bp) |
| *Mane-B\*007:03* | IPD0007057 | *Mamu-B\*007:04:01* (IPD0002243 - 1089 bp),<br>*Mafa-B\*007:08* (IPD0006970 - 1089 bp) |
| *Mane-B\*013:02* | IPD0007058 | *Mamu-B\*013:01* (IPD0002244 - 1089 bp),<br>*Mafa-B\*013:18* (IPD0007441 - 1089 bp) |
| *Mane-B\*014:02:02N* | IPD0007196 | *Mamu-B\*014:01* (IPD0002245 - 1089 bp),<br>*Mafa-B\*014:01* (IPD0002745 - 1089 bp) |
| *Mane-B\*027:03:02* | IPD0004174 | *Mafa-B\*044:06* (IPD0004326 - 1087 bp),<br>*Mamu-B\*027:02* (IPD0002261 - 1054 bp) |
| *Mane-B\*031:01* | IPD0006352 | *Mafa-B\*031:03* (IPD0007452 - 1095 bp) |
| *Mane-B\*045:03* | IPD0007065 | *Mamu-B\*045:08:01* (IPD0008784 - 1089 bp) |
| *Mane-B\*046:01:01* | IPD0004143 | *Mafa-B\*046:01:01* (IPD0002699 - 1080 bp) |
| *Mane-B\*051:07* | IPD0007212 | *Mafa-B\*051:04:01* (IPD0003827 - 425 bp) |
| *Mane-B\*055:01* | IPD0007214 | *Mafa-B\*055:01* (IPD0005568 - 659 bp) |
| *Mane-B\*060:02* | IPD0004151 | *Mafa-B\*060:01* (IPD0001682 - 1080 bp) |
| *Mane-B\*064:02* | IPD0007221 | *Mamu-B\*064:01* (IPD0002308 - 1077 bp),<br>*Mafa-B\*064:05* (IPD0007462 - 1077 bp) |
| *Mane-B\*065:01:01* | IPD0007223 | *Mafa-B\*065:05* (IPD0007463 - 1089 bp) |
| *Mane-B\*068:02:03* | IPD0004155 | *Mamu-B\*068:02:01* (IPD0006301 - 1089 bp),<br>*Mafa-B\*068:06:01* (IPD0006510 - 1089 bp) |
| *Mane-B\*068:07:01* | IPD0006370 | *Mafa-B\*068:03* (IPD0002713 - 1089 bp) |
| *Mane-B\*082:04* | IPD0007072 | *Mafa-B\*082:01:01* (IPD0003662 - 1107 bp) |
| *Mane-B\*089:02* | IPD0002914 | *Mamu-B\*089:01:02* (IPD0004042 - 822 bp) |
| *Mane-B\*089:03* | IPD0004167 | *Mafa-B\*089:01:02* (IPD0002039 - 1116 bp),<br>*Mamu-B\*089:01:02* (IPD0004042 - 822 bp) |
| *Mane-B\*105:01* | IPD0002765 | *Mafa-B\*105:01* (IPD0004004 - 1089 bp),<br>*Mamu-B\*105:01* (IPD0003536 - 649 bp) |
| *Mane-B\*105:03* | IPD0007088 | *Mamu-B\*105:01* (IPD0003536 - 649 bp) |
| *Mane-B\*136:01* | IPD0007202 | *Mafa-B\*136:02* (IPD0001651 - 1089 bp) |
| *Mane-B\*144:01* | IPD0004181 | *Mafa-B\*144:02* (IPD0002698 - 1089 bp) |
| *Mane-I\*01:01:01* | IPD0000633 | *Mafa-I\*01:11:01* (IPD0006272 - 1089 bp) |
| *Mane-I\*01:20:01* | IPD0007109 | *Mafa-I\*01:01:01* (IPD0000353 - 1089 bp) |
| *Mane-I\*01:22:01* | IPD0007176 | *Mamu-I\*01:06:01* (IPD0001264 - 1089 bp) |
| *Mane-I\*01:27* | IPD0007246 | *Mamu-I\*01:29* (IPD0007009 - 1089 bp),<br>*Mafa-I\*01:09* (IPD0002494 – 1089 bp) |
| *Mane-I\*01:36* | IPD0009089 | *Mamu-I\*01:30* (IPD0007011 - 1089 bp) |

Summary of 37 *Mane* full-length ORF sequences that are identical to previously described Mamu and/or Mafa nucleotide sequences

### Haplotype characterization

We used the full-length ORF sequence genotyping results to determine high-resolution haplotypes. The *Mane-A* and *Mane-B* regions were examined independently since these gene clusters are each separated by ~1Mb and recombination events are relatively common on a population basis. Haplotypes were characterized by comparing samples with identical groupings of alleles, particularly animals with known relationships. Pedigree information was available for most of the animals in these cohorts. Of the 194 animals examined, 133 were known to be directly related to at least one other animal sequenced in the PacBio experiments. For those animals that lacked pedigree information, relatedness could be inferred based upon the prevalence and sharing of specific haplotypes. Haplotype designations were assigned first for combinations of alleles observed together in related animals. We then looked across the full cohort to identify all animals expressing these initially defined haplotypes. For any animals with a single haplotype defined after this process, the remaining unassigned alleles were inferred to be on the alternate parental haplotype. To define and name the haplotypes, Mane-A and Mane-B sequences were first divided roughly into major and minor transcripts - sequences that averaged greater than 4% abundance relative to the total number of sequence reads identified per sample were denoted as major transcripts, and sequences that averaged lower than 2% abundance per sample were denoted as minor transcripts. In order to be uniform across macaque species, transcripts averaging in the intermediate range between 2% - 4% abundance per sample were denoted as major or minor largely in accordance with their designations in rhesus or cynomolgus macaques (Karl et al. 2013, 2017).

Of the major, highly expressed transcripts on each haplotype, one sequence was designated as the ‘diagnostic’ major allele, typically, the most abundant transcript on each haplotype. A few exceptions to this rule were made for sequences with putative biological significance based on studies in other macaque species, e.g., the B017 group of haplotypes. The initial four digits of each haplotype designation derive from the diagnostic transcript sequence (B004, B008, etc.). Haplotypes with different lineage groups of major alleles traveling with the diagnostic major are denoted with suffixes of lowercase letters. Thus, the B013a haplotype consists of a *Mane-B*013* diagnostic major transcript sequence accompanied by *Mane-B*014*, *Mane-B*041* and *Mane-B*178* transcripts while the B013b haplotype consists of a *Mane-B*013* diagnostic major transcript plus a *Mane-B*007* transcript. Finally, for the high-resolution haplotypes defined here, any sub-haplotypes differing from each other by any allelic variants across the haplotype are assigned a different Roman numeral suffix. For instance, the B013a.i sub-haplotype consists of the major alleles *Mane-B*013:01*, *Mane-B*014:01N*, *Mane-B*041:05*, and *Mane-B*178:01:02* while the B013a.ii sub-haplotype is defined by the major alleles *Mane-B*013:01*, *Mane-B*014:05*, *Mane-B*041:05*, and *Mane-B*178:01:02*. Therefore, this pair of sub-haplotypes differ only in the specific variants of *Mane-B*014* allele that were observed. Most of these major haplotype variants also differed in the minor MHC-I transcripts that were detected to be co-inherited, but that level of distinction was not considered when subdividing haplotypes with letter and/or Roman numeral suffixes. Figures 1a and 1b illustrate how haplotypes are assigned within related groups of animals.

**Fig. 1a.**
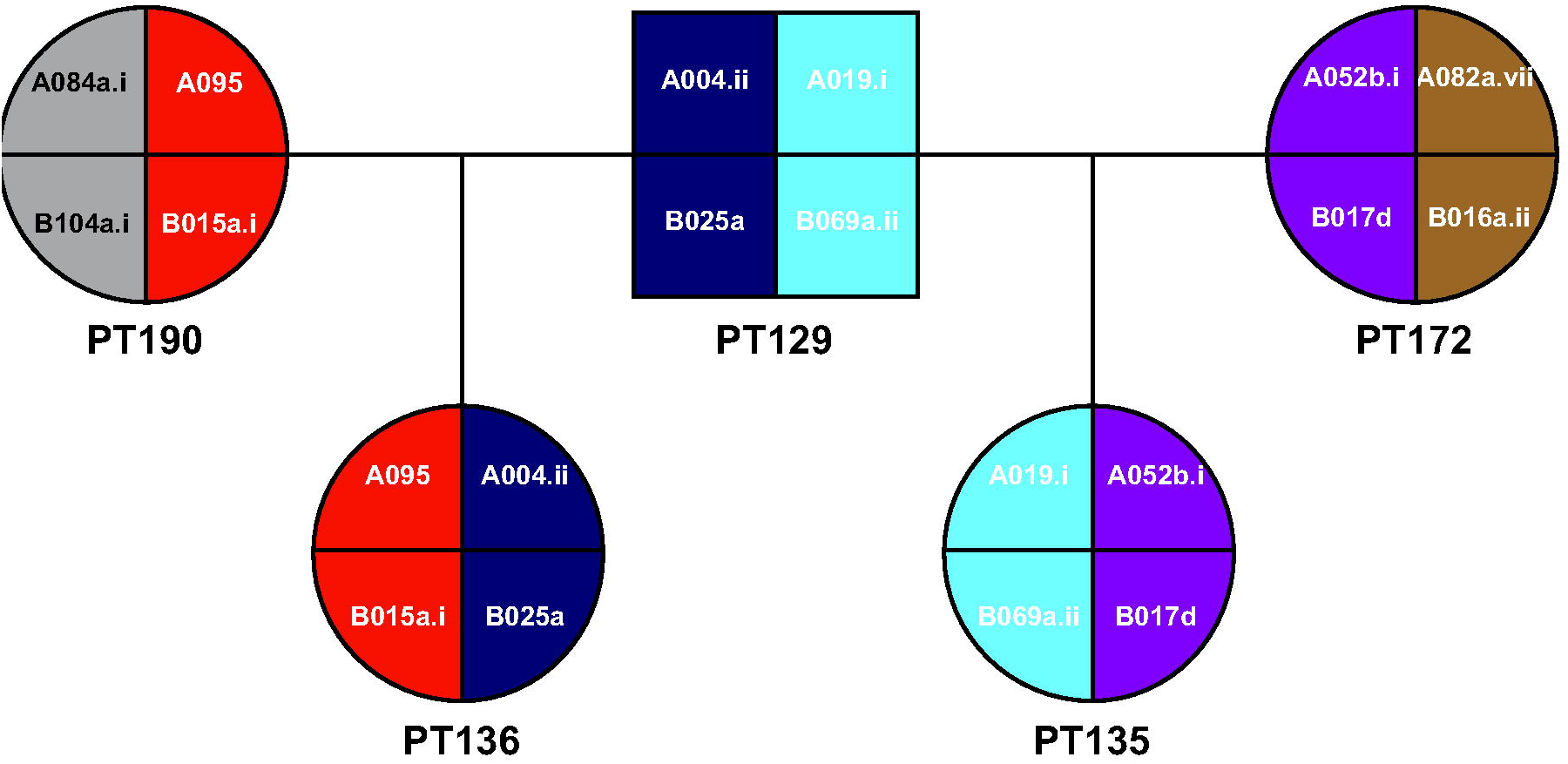
Pedigree describing haplotype inheritance. Sample pedigree information visually showing the passage of MHC-I haplotypes from sires (PT129) and dams (PT172 and PT190) to their respective progeny (PT135 and PT136)

**Fig. 1b.**
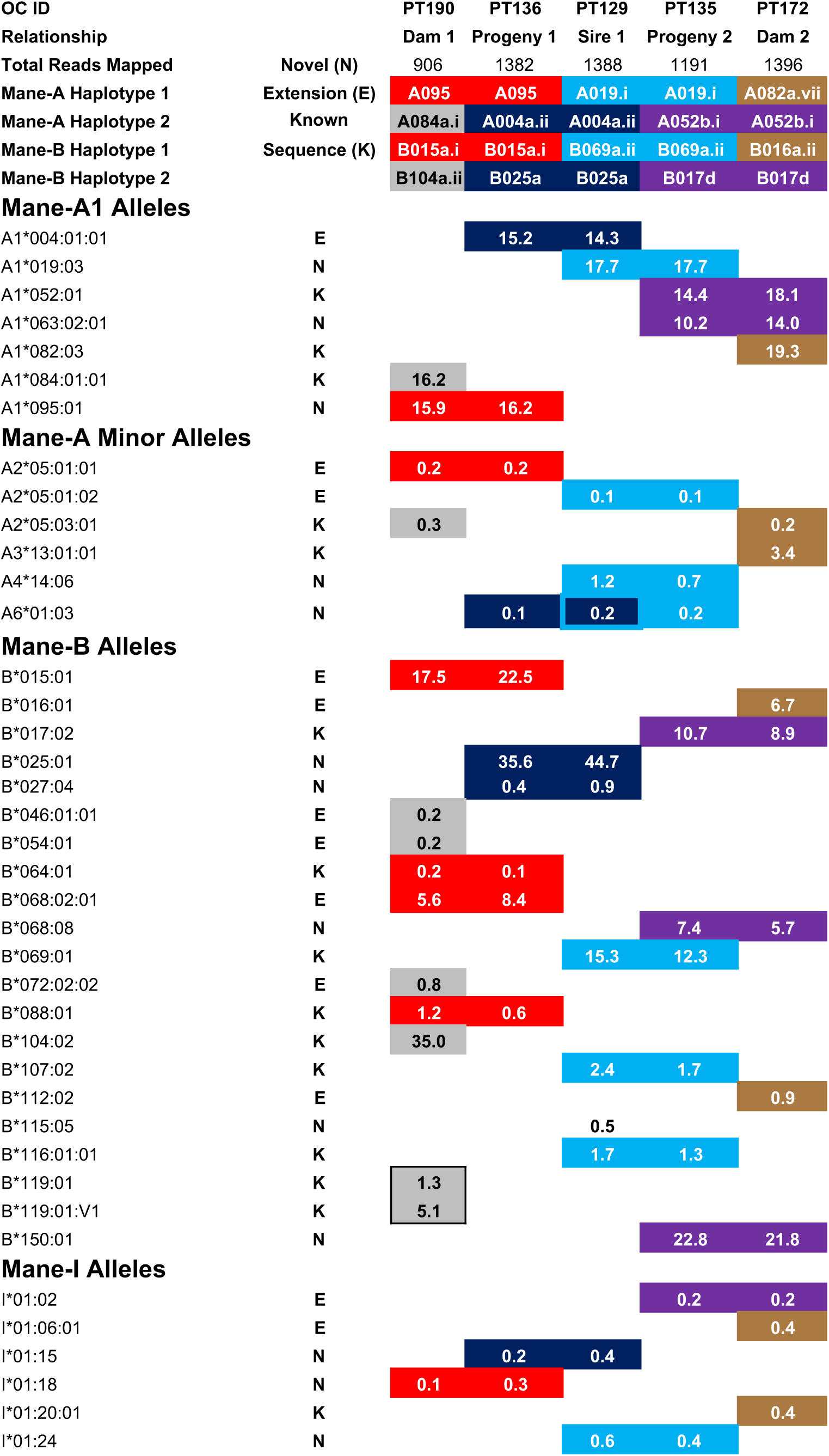
Inheritance of alleles encompassing each haplotype. A subsequent representation of the pedigree information showing the passage of the common transcripts seen in each haplotype to the various offspring. The alleles are also represented as percent of the total sequences observed in each animal providing a way to distinguish between major and minor transcript sequences in the haplotype

In total, we characterized 192 *Mane* class I haplotypes; 86 of these were *Mane-A* haplotypes and 106 of these were *Mane-B* haplotypes. The definitions of the *Mane-A* and *Mane-B* haplotypes are shown in Figures 3 and 4 respectively. This distribution is consistent with previous macaque data sets (Karl et al. 2013, 2017) where MHC haplotypes of the B genes are more numerous; presumably this is due to the additional rounds of complex duplication events that macaque B regions have experienced. Of the *Mane-A* haplotypes, A052a.i was the most common, being observed in 13.4% of the 388 total chromosomes. Likewise, B118d was the most common *Mane-B* haplotype; it was present in 7.5% of the total chromosomes examined.

**Fig. 2a.**
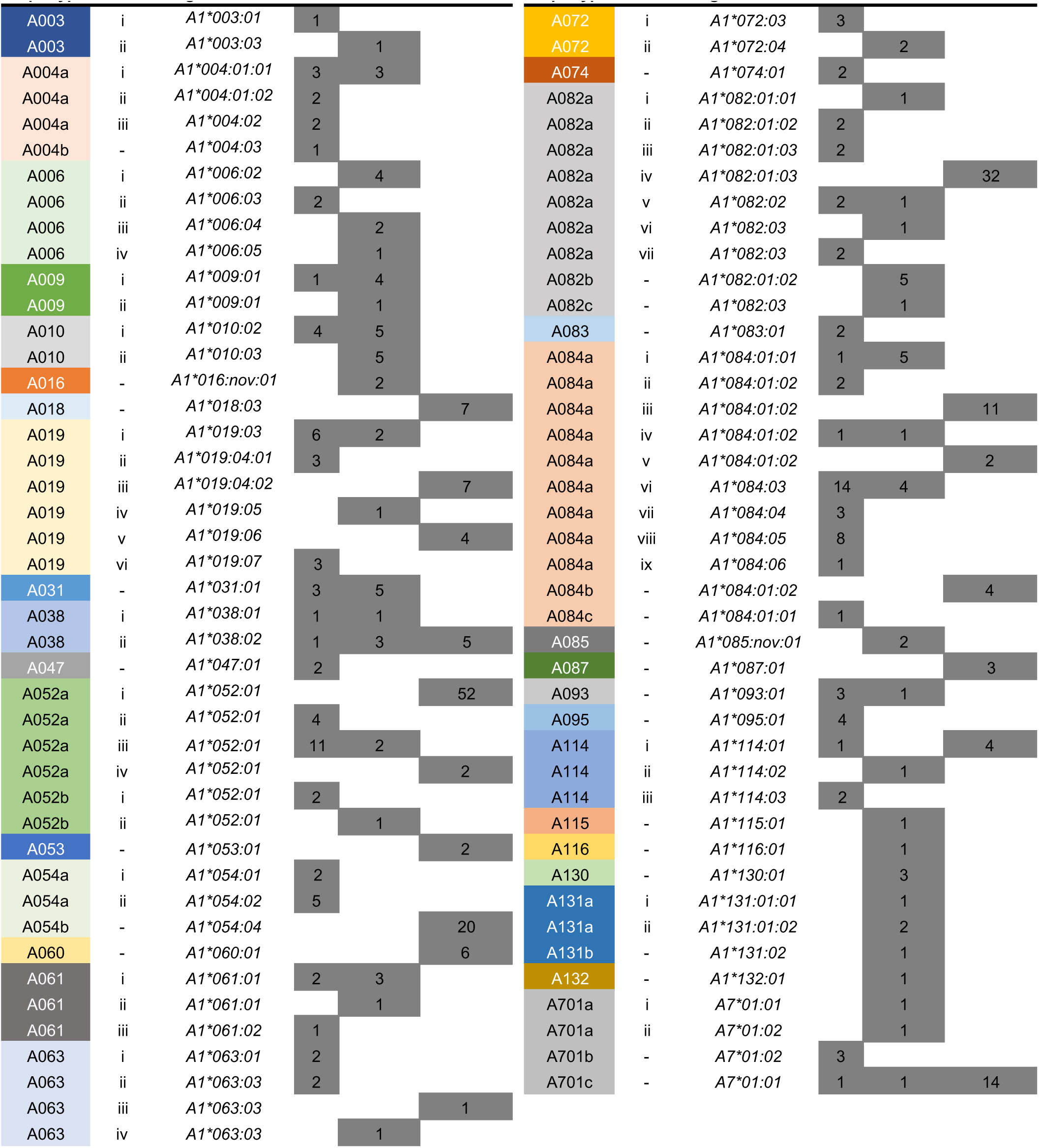
Distribution of *Mane-A* haplotypes across research institutions. Summary of the distribution of Mane-A haplotypes across the three cohorts in the study. Each haplotype and variant is represented by the diagnostic transcript for which it is named with the number of chromosomes each haplotype was observed in indicated

**Fig. 2b.**
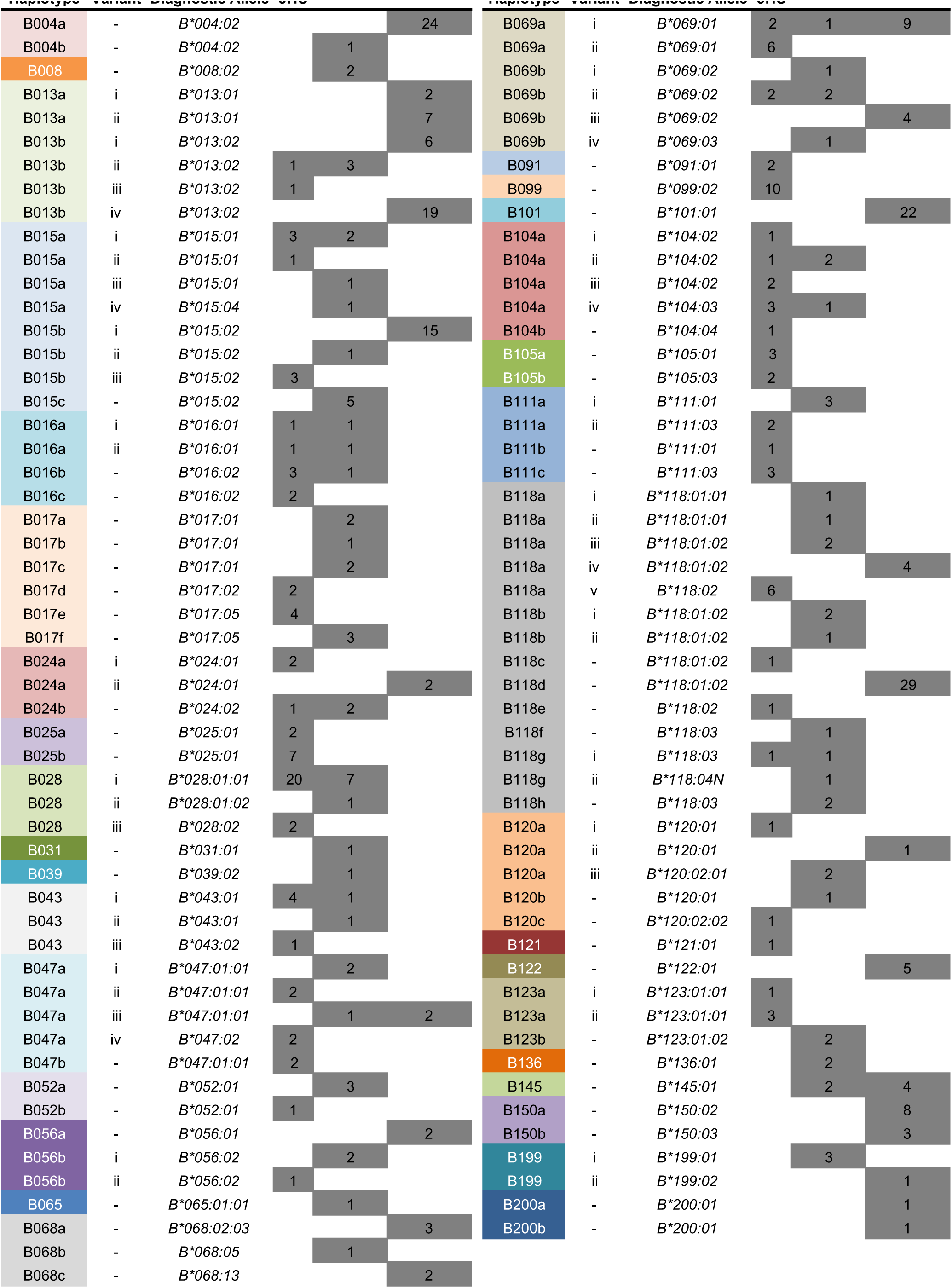
Distribution of *Mane-B* haplotypes across research institutions. Summary of the distribution of Mane-B haplotypes across the three cohorts in the study. Each haplotype and variant is represented by the diagnostic transcript for which it is named with the number of chromosomes each haplotype was observed in indicated

**Fig. 3.**
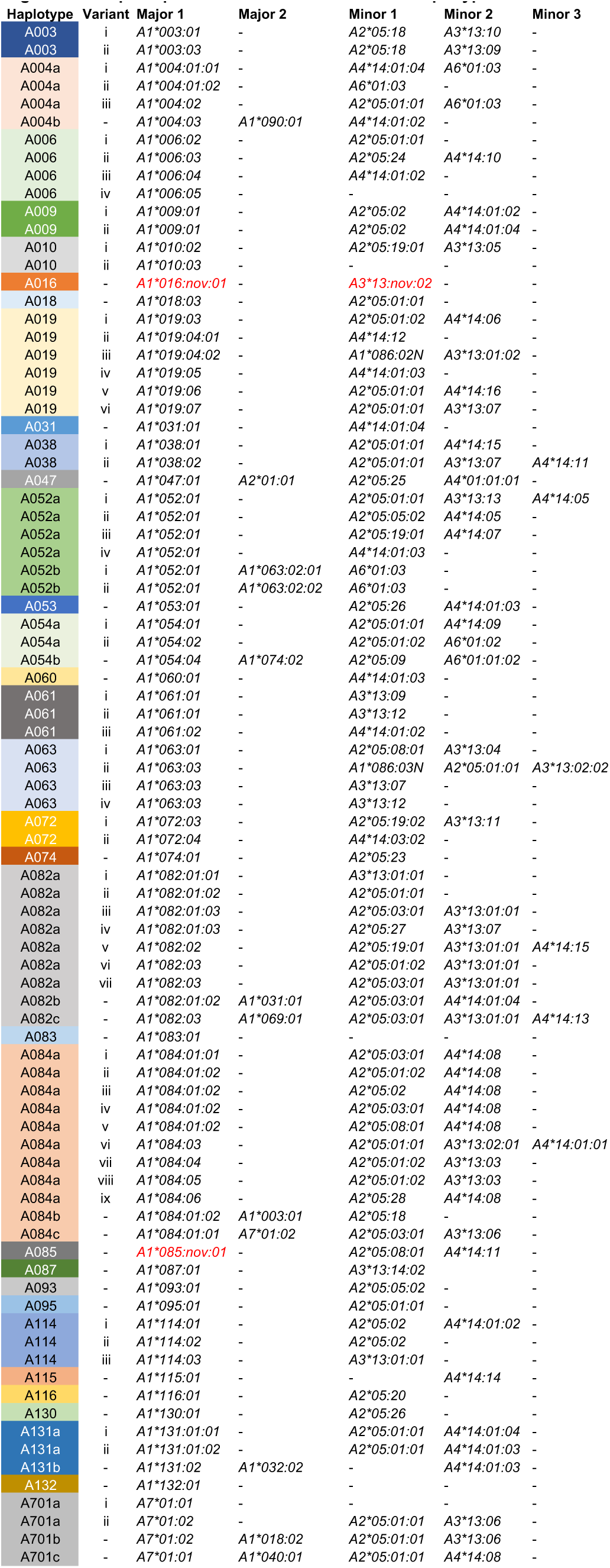
Transcript sequences associate with *Mane-A* haplotypes. A description of the sequences that define each *Mane-A* haplotype. The haplotypes are named for the first major transcript described and contain other major and minor alleles that are passed onto offspring together. Major transcripts were defined as having over 4% of the total reads associated with each animal and minor alleles had less than 2% of the total reads. Sequences with an abundance between 2-4% were defined as major or minor in accordance with published alleles from rhesus or cynomolgus macaques (Karl et al. 2013, Karl et al. 2017)

**Fig. 4.**
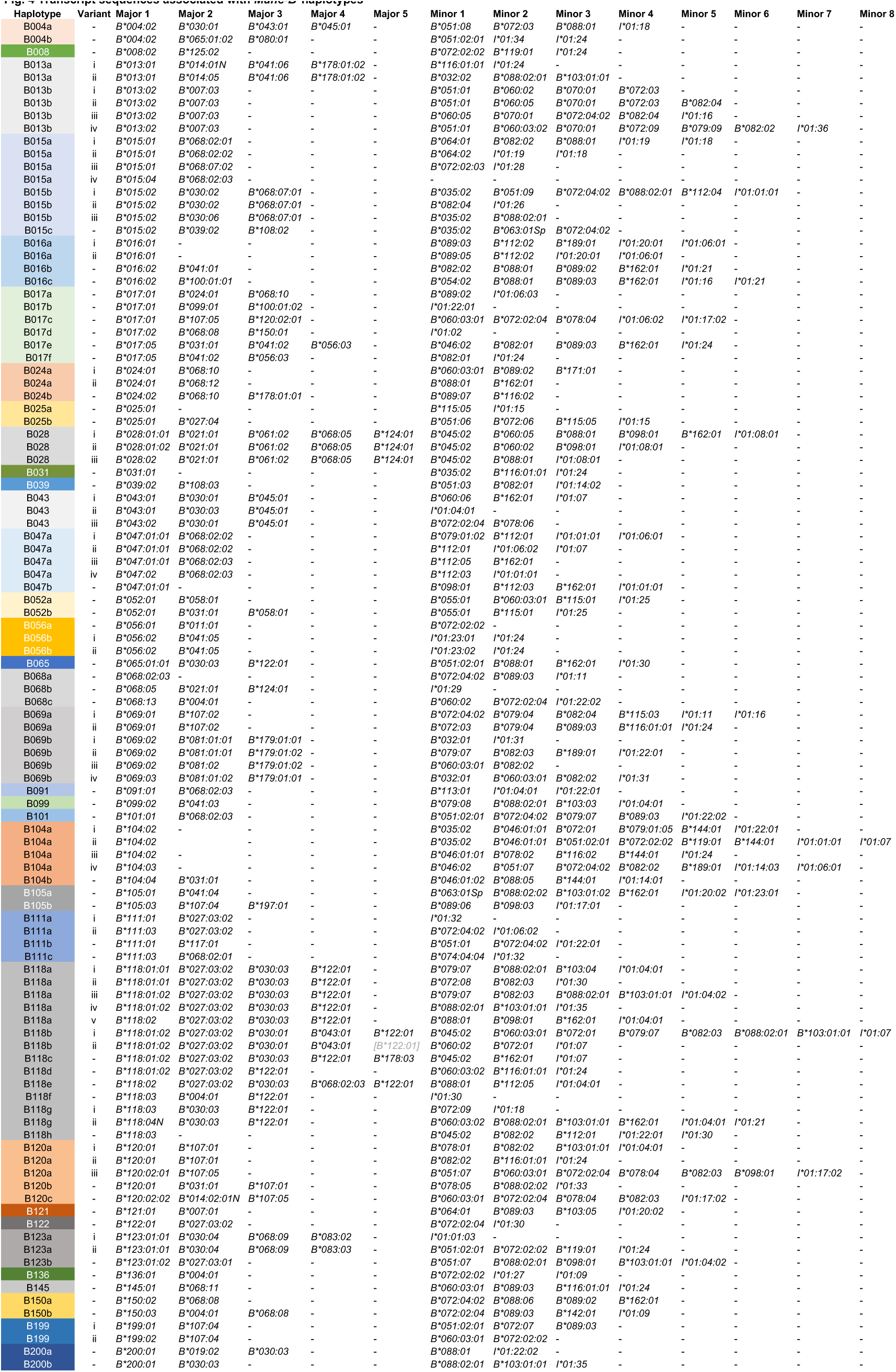
Transcript sequences associated with *Mane-B* haplotypes. A description of the sequences that define each *Mane-B* haplotype. The haplotypes are named for the first major transcript described and contain other major and minor alleles that are passed onto offspring together. Major transcripts were defined as having over 4% of the total reads associated with each animal and minor alleles had less than 2% of the total reads. Sequences with an abundance between 2-4% were defined as major or minor in accordance with published alleles from rhesus or cynomolgus macaques (Karl et al. 2013, Karl et al. 2017)

### Haplotype diversity among breeding centers

The primary goal of these studies was to maximize the discovery of novel *Mane-A* and *Mane-B* sequence variants and to characterize additional high-resolution *Mane-A* and *Mane-B* haplotypes. With this goal in mind, cDNA from animals at the JHU and WaNPRC breeding centers was prescreened by Illumina MiSeq analyses in order to enrich these PacBio cohorts with individuals expected to carry novel class I alleles/haplotypes. A subset of MHC-identical animals from common sires and dams in these breeding groups were excluded in order to limit redundancy in the PacBio analyses. In contrast, all cDNA samples from two consecutive years of health screening for the pig-tailed macaques in the University of Melbourne and Monash University breeding program were analyzed sequentially without restricting the number of offspring from common sires and dams that were evaluated. Compared to the other breeding centers, this methodological difference resulted in apparently lower MHC haplotype diversity for animals in the Australian cohort where haplotypes A052a.i and A082a.iv together made up almost half of the *Mane-A* haplotype distribution (Figure 2a). The distribution of *Mane-B* haplotypes was slightly more diverse, but four haplotypes, B118d, B004a, B101 and B013b.iv accounted for just over 50% of the total haplotypes observed in this population (Figure 2b).

The majority of *Mane-A* and *Mane-B* haplotypes defined in this study were specific to a single institution. However, multiple haplotypes were present across more than one cohort, as 16% of the *Mane-A* haplotypes and 14% of the *Mane-B* haplotypes were shared between at least two breeding centers. Most of the shared *Mane-A* and *Mane-B* haplotypes were observed between JHU and WaNPRC which was expected because a subset of the founding pig-tailed macaques for the JHU breeding program were obtained from WaNPRC. In contrast, only six *Mane-A* and *Mane-B* haplotypes observed in individuals from the Australian cohort were also shared with animals evaluated from JHU and WaNPRC. Additional studies will be required in order to determine whether these differences reflect distinct geographic origins for the pig-tailed macaques at each of these breeding centers.

### Advantages of using full-length cDNA sequencing over short genotyping amplicons

In understudied populations of macaques and other NHPs, PacBio full-length sequencing provides multiple advantages over use of short amplicons that have been used routinely for MHC genotyping analyses over the past several years (Karl et al. 2013, 2017). For one, pig-tailed macaques and other understudied NHP populations have been historically used less often in infectious disease research compared to rhesus macaques, so the MHC region of these species is generally less well characterized. PacBio full-length sequencing provides an opportunity to not only discover novel allelic variants, but also extend partial sequences to include complete ORFs. It also provides us with improved resolution between related sequences, allowing for alleles to be defined to the level of synonymous variation in the genetic sequence. By defining these transcript sequences with higher specificity, it becomes possible to study whether functional differences arise due to subtle variations in the genetic sequence. Another advantage of defining full-length ORF sequences is in regards to restriction of infectious diseases. Previous studies have identified multiple MHC-I variants that bind to specific epitopes of SIV (Gooneratne et al. 2014). In this study, we identified multiple closely related allelic variants of these MHC-I gene products that have been documented to restrict SIV epitopes. Because of the subtle, but possibly important differences in these newly discovered allelic variants, they may present distinct target epitopes from SIV/HIV and other infectious agents. Further research is needed to characterize immune responses associated with these allelic variants, but their discovery may be an important piece of information for the use of an effective long-term immune response against HIV or novel vaccine development.

### Matching MHC identical animals to prevent GvHD

An additional advantage of using full-length sequencing over shorter amplicons involves transplant research. Full-length allele discovery defines transcript sequences to the level of nonsynonymous and synonymous polymorphisms in all coding regions of the gene. This specificity is of great use in transplant research where having MHC identical donors and recipients is necessary to reduce the chances of GvHD, in which the host cells recognize the newly transplanted cells as a foreign antigen and develop an immune response (Anasetti et al. 1990; Ayala García et al. 2012). It is also advantageous to match MHC or human leukocyte antigen (HLA) donors and recipients in terms of stem cell transplants. A recent study described the use of induced pluripotent stems cells in the brain to reduce the effects of Parkinson’s disease (PD) by regenerating and increasing the survival of dopamine releasing neurons. The MHC identical NHPs used in this study showed a reduced immune response from microglia and lymphocytes to the graft (Morizane et al. 2017). By continuing to describe MHC-I transcripts and haplotypes to specific levels of allelic variation, animals can be matched to minor allelic variation and GvHD-like diseases can be reduced in studies. It also provides a means for future studies to be undertaken in the hopes that these resources and therapies can provide a means for treating difficult neurological disorders.

However, with this specific level of definition, a paradoxical situation may arise in which as we identify more allelic variants, it may become difficult to find donors and recipients that have genetically matching MHC. As a counter argument, there may a difference between being genetically identical and being functionally identical in terms of the host response in GvHD. There is a possibility that, although there are amino acid differences between two variants of the same MHC-I sequence, there may not be significant differences in their functionality and phenotypic appearance to host cells. The discovery of novel transcripts and further haplotype definition will enhance the specificity to which the MHC-I proteins are defined, but further research is necessary to define whether this level of definition gives rise to phenotypic differences noticeable to host immune cells.

In contrast, GvHD may be a beneficial component to potential HIV cure strategies. Timothy Ray Brown, better known as the “Berlin Patient” was “cured” of HIV when he received an allogeneic hematopoietic stem cell transplant from a homozygous CCR5Δ32 donor after intensive chemotherapy. Almost ten years after his transplant, he maintains undetectable levels of HIV DNA and RNA without antiretroviral therapy (ART) (Yukl et al. 2013). It has been postulated that one of the mechanisms involved in treating, and eventually controlling his infection with HIV was essentially a graft-versus-host effect that targets latently infected cells, thus depleting the HIV reservoir (Mavigner et al. 2014; Zou et al. 2013). With our high-resolution MHC genotyping approach, it becomes possible to design future studies with more control over the effects due to transplantation. Due to their infectability with HIV and progression to AIDS (Baroncelli et al. 2008, Hatziioannou et al. 2014), pig-tailed macaques provide an important model to test this HIV cure strategy.

### Pig-tailed macaques and their use in infectious disease research

Pig-tailed macaques are important models for infectious disease research, and especially for research involving HIV, SIV and AIDS-like diseases. Pig-tailed macaques express a variant of the TRIM5α protein that allows them to be infected with minimally modified forms of HIV-2 and multiple forms of SIV (Brennan et al. 2007; Hatziioannou et al. 2009, 2014; Igarashi et al. 2007; Kirmaier et al. 2010). Because of this, pig-tailed macaques provide a more accurate representation of the course of HIV infection in humans and progression to AIDS-like diseases (Baroncelli et al. 2008, Hatziioannou et al. 2014).

Certain MHC haplotypes provide protection against SIV and may be able to slow progression of SIV into AIDS-like diseases in macaques. Notably, in pig-tailed macaques, *Mane-A1*084:01*, previously named *Mane-A*10*, restricts the Gag KP9 and several additional epitopes of SIV, thus slowing SIV viral escape in T cell lymphocytes (Gooneratne et al. 2014). SIV viral loads in the plasma are also shown to be significantly reduced in macaques expressing *Mane-A1*084* compared to macaques without it (Smith et al. 2005a, b). In the current study, we identified two additional variants of *Mane-A1*084* lineage (*Mane-A1*084:05* and *Mane-A1*084:06*) that differed from *Mane-A1*084:01* by one and two nonsynonymous substitutions respectively. While there is uncertainty whether or not all seven members of the *Mane-A1*084* lineage provide the same level of protection, the fact that they differ from the reference sequence by such a small number of amino acid substitutions leads us to believe that a similar level of protection is possible. Further studies are required in order to make conclusions on the protectiveness of these alleles. If our novel sequences do provide the same level of protection, there is incentive for researchers to use animals containing these alleles for further studying HIV progression and escape in pig-tailed macaques.

Other species of macaques have documented alleles that are shown to restrict progression of SIV to AIDS like disease. Notably in rhesus macaques, both *Mamu-B*008:01* and *Mamu-B*017:01:01* have been shown to control SIV replication and progression to disease; *Mamu-B*008:01* restricts the Vif RL8, Vif RL9, and Nef RL10 epitopes of SIV and *Mamu-B*017:01:01* restricts the Nef IW9 epitope (Loffredo et al. 2007; Martins et al. 2015; O’Connor et al. 2003; Yant et al. 2006). In our studies, we identified one novel variant of both *Mane-B*017:01* and *Mane-B*008:01*. Both the novel variant of *Mane-B*008:01* and *Mane-B*017:01* discovered in these cohorts differ from the previously characterized *Mamu* versions by two nonsynonymous amino acid substitutions. Our novel variant of *Mane-B*008:01* was seen in four animals from the WaNPRC at relatively high numbers of the total reads from each of the four animals. This observation is consistent with high levels of transcription for the specific sequence, as has been observed for *Mamu-B*008:01* in rhesus macaques. The novel *Mane-B*017:01* variant was seen in six animals coming from both the JHU colony and the WaNPRC, again in relatively high read numbers. Since these novel variants are relatively common in these cohorts, they may be useful models as to test whether the newly characterized MHC transcript sequences show the same protective effects as those observed in their *Mamu* counterparts (Gooneratne et al. 2014; Loffredo et al. 2007; Martins et al. 2015; O’Connor et al. 2003; Yant et al. 2006). Previous studies have described specific polymorphisms in multiple SIV genes that are linked to related alleles and haplotypes (Gooneratne et al. 2014). However, subsequent studies are required to show a correlation between these newly discovered alleles and control of SIV replication as has been seen in rhesus macaques.

The increased allelic resolution and discovery provided by PacBio CCS adds to our knowledge of pig-tailed macaques while also representing an important step for advancing their use in biomedical research. As an important model for studying HIV infection and progression to AIDS-like disease, the use of full-length sequencing technology and the discovery of novel allelic variants that may be protective against infection promote the use of the pig-tailed macaque model for infectious diseases.

## Acknowledgements and Funding Sources

The authors gratefully acknowledge the support of Suzanne Queen at Johns Hopkins University, Rita Cervera Juanes and Betsy Ferguson at the Oregon Health & Science University, and Thakshila Amarasena and Stephen Kent at the University of Melbourne for their assistance in the collection and shipping of pig-tailed macaque RNA and cDNA. We also would like to thank Nel Otting, Natasja de Groot, and Ronald Bontrop at the Biomedical Primate Research Centre for their assistance in submitting and naming of novel sequences in the Immuno Polymorphism Database for the Major Histocompatibility Complex genes of Non-Human Primate.

This research was supported by contracts HHSN272201600007C and HHSN272201100013C from the National Institute of Allergy and Infectious Diseases and the National Institutes of Health, and was conducted at a facility constructed with support from the Research Facilities Improvement Program (RR15459-01, RR20141-01).

**ESM 1** PacBio full-length amplicon primer design. Unique barcodes are added to the 5’ and 3’ end of each transcript and hairpin loops are created during library preparation for sequencing on the PacBio RS II instrument

**ESM 2** Full genotyping table including animals from all three cohorts showing the number of reads for each transcript sequence seen in the animals. Alleles are color coded according to the haplotype that they are passed along with, and haplotypes passed on from sires or dams to offspring are denoted with colored text

